# Diversity Converges During Community Assembly in Methanogenic Granules, Suggesting a Biofilm Life-Cycle

**DOI:** 10.1101/484642

**Authors:** Anna Christine Trego, Cristina Morabito, Simon Mills, Stephanie Connelly, Isabelle Bourven, Giles Guibaud, Christopher Quince, Umer Zeeshan Ijaz, Gavin Collins

**Affiliations:** School of Natural Sciences, National University of Ireland Galway, University Road, Galway, H91 TK33, Ireland; School of Engineering, University of Glasgow, Oakfield Avenue, Glasgow G128LT, United Kingdom; Groupement de Recherche Eau Sol Environnement, Faculté des Sciences Techniques, Université de Limoges, 123 Avenue Albert Thomas, 87060 Limoges Cedex, France; Warwick Medical School, University of Warwick, Gibbet Hill Road, Coventry CV4 7AL, United Kingdom

## Abstract

Anaerobic biological decomposition of organic matter is ubiquitous in Nature wherever anaerobic environments prevail, and is catalysed by hydrolytic, fermentative, acetogenic, methanogenic, and various other groups, including syntrophic bacteria. It is also harnessed in innovative ways in engineered systems that may rely on small (0.1-4.0 mm), spherical, anaerobic granules, which we have found to be highly-replicated, whole-ecosystems harbouring the entire community necessary to mineralise complex organics. We hypothesised distinct granule sizes correspond to stages in a biofilm life-cycle, in which small granules are ‘young’ and larger ones are ‘old’. Here, granules were separated into 10 size fractions used for physico-chemical and ecological characterisation. Gradients of volatile solids, density, settleability, biofilm morphology, methanogenic activity, and EPS profiles were observed across size fractions. Sequencing of 16S rRNA genes indicated linear convergence of diversity during community assembly as granules increased in size. A total of 155 discriminant OTUs were identified, and correlated strongly with physico-chemical parameters. Community assembly across sizes was influenced by a niche effect, whereby *Euryarchaeota* dominated a core microbiome presumably as granules became more anaerobic. The findings indicate opportunities for precision management of environmental biotechnologies, and the potential of aggregates as playgrounds to study assembly and succession in whole microbiomes.

## MAIN

### Introduction

Global carbon cycling comprises not only primary production by photo- and chemosynthesis on one hand, and carbon consumption by respiration on the other, but also relies fundamentally on the anaerobic, biological decomposition of organic matter wherever suitable environments prevail, such as in saturated wetlands; coastal, lake and marine sediments; and ruminant and termite guts, among many others [1–3]. Anaerobic digestion (AD) is a natural process mediated by the collective, sequential and cooperative action of several trophic groups of microorganisms, including hydrolytic bacteria, fermenters, organic-acid-oxidisers, and – finally – evolutionarily ancient, methanogenic archaea feeding on a narrow range of substrates including acetate, methanol and H_2_/CO_2_. Methanogenic consortia of this type comprise complex microbial communities of bacteria and archaea, and may include a diversity of syntrophic species, which operate within extremely narrow thermodynamic windows of profitability [4], as well as diversionary organisms – such as sulfate-reducing bacteria – competing with others for energy.

The AD process is widely harnessed for wastewater treatment in engineered, digester systems [5–7]. Many such digesters apply planktonic, suspended microbial cells to stabilise, and valorize, high-solids wastes, such as manures or crop-residues. Those are typically low-rate systems – applied under relatively low volumetric loading and, hence, across long hydraulic retention times so as to prevent washout from the digester of active cells. However, a distinct family of anaerobic digesters includes ‘retained-biomass’ systems based on the use of self-immobilised, biofilm granules, which settle quickly and, thus, avoid washout even from digesters applied under conditions of high volumetric loading and short hydraulic retention [8]. In this way, the use of anaerobic granules allows decoupling of hydraulic and biomass retention in anaerobic digesters, and such biofilms may be retained in AD systems for long periods of at least weeks or months.

Anaerobic granules, which were first discovered in the 1970s [9,10] and have since revolutionised the treatment of industrial wastewaters [11], are spherical biofilms of approximately 0.2-5 mm in diameter, each comprising a diverse microbial consortium, and each theoretically providing the entire microbial community necessary for complete mineralisation of complex organic feedstocks and wastes. The granular structure supports efficient transfer of substrates between trophic groups, as well as providing protection from toxins and environmental perturbations. Hulshoff Pol *et al.* [12] provided a pivotal review comparing mechanisms, and proposed a diversity of theories – based on ecology, physico-chemical features, and thermodynamics – to explain the granulation process, which all generally agreed that inert carriers likely play a critical role in granule formation, and that the early stages of granule development likely mimic the classical models of cell attachment and biofilm formation on solid surfaces [13,14]. Most authors also agree that *Methanosaeta concilii*, a methanogen with high acetate affinity, plays a key granulation role in providing a core of tangled filaments around which other cells aggregate. Whilst several studies focused on the earliest stages of biofilm development, very few, however, studied growth and maturation of granules, and the microbiome, over time [12].

Indeed, many studies have described particle formation from diverse microbial communities, such as the bacterial colonization in the ocean of particulate organic matter [15], but the mechanisms by which complex communities assemble to form stable biofilms are still only poorly understood. Others describe the interaction of neutral and species-sorting processes, and the roles of generalists and specialists in community assembly [16].

Whilst full-scale anaerobic digesters (with volumes of typically up to several hundred cubic metres) contain millions of single-granule ecosystems, not all granules are the same. Ahn [17] found the typical range of granule diameters to be 0.1-5 mm, and in a characterisation by Diaz *et al.* [18] different granule morphologies were observed from single digesters. They separated granules by colour, and differences were observed in granule size and structure. Nonetheless, our recent research has indicated that granules from within distinct size ranges harbour statistically identical microbiomes [19], even though the community structure in granules of different sizes may differ. Each individual granule therefore represents a perfectly parameterised, whole biofilm ecosystem. Thus – and rather uniquely in either Nature or the built environment – anaerobic granules may well be considered as distinct, replicated, whole microbial communities.

This may pose an opportunity to test questions about community assembly; growth patterns; and drivers of community structure and diversity in complex biofilms – including as encouraged by Rillig *et al.* [20], who proposed soil aggregates as massively concurrent evolutionary incubators. The utility of such highly-replicated, whole-microbial-communities to study microbial evolution should be explored. For example, the extent to which ecological theories hold across the expansion of, and succession in, the microbial community in granules should provide useful information for the management of environmental biotechnologies. Moreover, whether the granulation process is cyclical, and follows a predictable life cycle, is an important question for both environmental engineering and microbial ecology.

We hypothesise that differently sized granules represent different stages of biofilm development and that granules taken from a single digester at a single point in time, having survived the same environmental conditions, may, in fact, represent different stages of growth over a biofilm life-cycle where the smallest granules are at the earliest stages of formation and the largest granules are the oldest and most mature. We attempted an intensive characterisation of anaerobic granules from a full-scale digester across multiple, discrete size fractions, to characterise morphological, physico-chemical, physiological and ecological differences across a set of highly-resolved granule size fractions. This provides an interesting new perspective for Microbial Ecology with respect to community assembly, biofilm development, and the drivers of microbial diversity in a controlled system underpinned by collections of single, whole-ecosystem aggregates.

### Materials and Methods

#### Source of biomass and size fractionation

Anaerobic sludge granules were obtained from a full-scale, mesophilic (37°C) upflow anaerobic sludge bed (UASB) digester treating potato-processing wastewater in the Netherlands. The sludge was size-separated into ten discrete size fractions (A-J) by passing the granules through a series of stainless-steel sieves. Each fraction was suspended in 1X phosphate buffered saline (PBS; Fisher Scientific, Geel, Belgium), sparged with N_2_ gas, and allowed to settle for 1 h before determining the settled volume.

#### Total, and volatile, solids; and sludge settling velocity, and density

The total solids (TS) and volatile solids (VS) concentrations of granules from each size fraction were determined using the standard loss-on-ignition technique [21]. The settling velocity, and density, of granules (n=10) from each size fraction, A-J, was determined. A 1-m long, clear, acrylic tube, fitted with a stopper at one end was fastened vertically and filled with deionised water. Two markings were made on the outside of the tube, at 0.3 m and 0.6 m from the top, and the water temperature was recorded. The diameter of individual granules was measured using electronic digital calipers. Granules were individually dropped into the column of water, and the time (seconds) required for each granule to travel 0.3 m (the distance between the two markings) was measured. The settling velocity was the distance (0.3 m) divided by settling time. Stokes’ Law was then applied to determine granule density.

#### Scanning electron microscopy (SEM)

Three granules from each size fraction were randomly selected for SEM imaging. Granules were placed in clean, individual 1.5-ml microcentrifuge tubes and covered with 2.5% (w/v) glutaraldehyde in 0.5 M cacodylate buffer (pH 7.2). The tubes were inverted gently and incubated overnight at 4°C. The supernatant was removed and granules were washed three times in 1X PBS, before being dehydrated by passing through a series of 10-min ethanol washes using 50%, 70% and 90% ethanol. Dehydrated granules were placed on carbon tabs, which were then fastened to aluminum stubs. An aliquot of 25 μl hexamethyldisilazane (HMDS) was placed on each granule, under a fume hood, and allowed to dry overnight. Specimens were gold-sputtered and imaged in a scanning electron microscope (Hitachi S-2600, Mountain View, CA, USA).

#### Extraction and characterisation of extracellular polymeric substances (EPS)

Loosely bound (LB) and tightly bound (TB) EPS was extracted in duplicate from sludge from each size fraction using the cation exchange resin (CER) technique [22,23]. Colorimetric assays were used to investigate the biochemical composition of the EPS using a spectral photometer (Cadas 50 S, Dr Lange, Berlin, Germany). Concentrations of proteins and humic-like substances (HLS) were determined and corrected [24,25]. Bovine serium albumin (96%, Sigma-Aldrich, St. Louis, Missouri, USA) was used as a standard for proteins and humic acids (Sigma-Aldrich, St. Louis, Missouri, USA) as the standard for HLS. Polysaccharides were measured following DuBois *et al.* [26], using a glucose standard.

#### Specific methanogenic activity (SMA)

An SMA buffer solution was prepared in a round-bottom flask by combining 0.4 ml 0.0001% (w/v) resazurin (Fisher Scientific, Geel, Belgium), 0.56 g cysteine hydrochloride monohydrate and enough distilled water (dH2O) to bring the volume to 700 ml. The pH was adjusted to 7.0-7.1 by dropwise addition of 8 M NaOH, and the final volume was adjusted to 1 L. The solution was boiled until clear, and immediately sealed and cooled on ice with constant N_2_ sparging until at 50°C when 3.05 g sodium bicarbonate were added before sealing the flask. The SMA buffer was added with sludge granules to 60-ml, glass bottles to give a final volume of buffer and granules of 10 ml, and a final VS concentration of 4 g L^−1^. The bottles were sealed and N_2_-flushed before acclimatisation at 37°C for 48 h. Aliquots of 0.1 ml soluble substrates were then added to separate, respective bottles to give final concentrations of 30 mM acetate, 15 mM butyrate or 30 mM propionate. No-substrate controls measured background activity. To test for autotrophic methanogenesis, H_2_/CO_2_ (80:20, v/v) was added at 1 bar for 20 s. N_2_/CO_2_ (80:20, v/v) was used to control H_2_-fed assays. Headspace biogas pressure was measured as millivolts (mv), using a handheld pressure transducer (CentrePoint Electronics, Galway, Ireland), and converted to biogas volume (ml) using a headspace correction factor [27,28]. Gas chromatography (CP-3800, VARIAN Inc., Walnut Creek, CA) was used to determine the methane concentration (%) in the biogas, and the accumulation rate was plotted. The precise, *in situ* concentration of VS in each bottle was determined by drying and burning, as before. In the case of soluble substrates, SMAs were determined under STP conditions as the daily rate of methane production as a function of the total VS. SMA for gaseous substrates were calculated using a similar approach; however, the rate was computed using the reaction stoichiometry of 4:1 molar of H_2_ consumption to methane production.

#### DNA Extraction

A mass of 0.1 g wet sludge from each of the size fractions was weighed into respective, sterile tubes in triplicate. DNA was extracted on ice following the DNA/RNA co-extraction method described by Griffiths *et al.* [29], which is based on bead beating in 5% (w/v) cetyl trimethylammonium bromide (CTAB) extraction buffer, followed by phenol-chloroform extraction. Integrity of nucleic acids was assessed using a nanodrop (Thermo Fisher Scientific, Waltham, MA, USA). Concentrations were determined using a Qubit fluorometer (Invitrogen, Carlsbad, CA, USA) and normalised to 5 ng DNA μl^−1^ before storage at −80°C.

#### High-throughput DNA Sequencing, Bioinformatics, and Statistical Analysis

Amplification of the 16S rRNA gene sequences was performed by The Foundation for the Promotion of Health and Biomedical Research of Valencia Region, FISABIO (Valencia, Spain) using the universal bacterial and archaeal primer set: forward primer 515F and reverse primer 806R. The resulting amplicon library of short inserts was sequenced on the Illumina MiSeq platform. Abundance tables were generated by constructing OTUs (as a proxy for species). Statistical analyses were performed in R using the combined data generated from the bioinformatics as well as meta data associated with the study. Details are described further in Supplementary Methods.

### Results

#### Distribution of granule sizes

A normal distribution of granule sizes was observed with a majority (>75%) of the sludge volume comprising medium-range granules (fractions D – G; Fig. 1). No granules >4 mm were found. The VS proportion of TS was relatively high (average, 91.8%; σ^2^, 0.88) in granules from medium and larger fractions (D – J) but lower in smaller granules (86.2%, 70.4% and 89.0% in fractions A, B and C, respectively).

**Figure 1.**
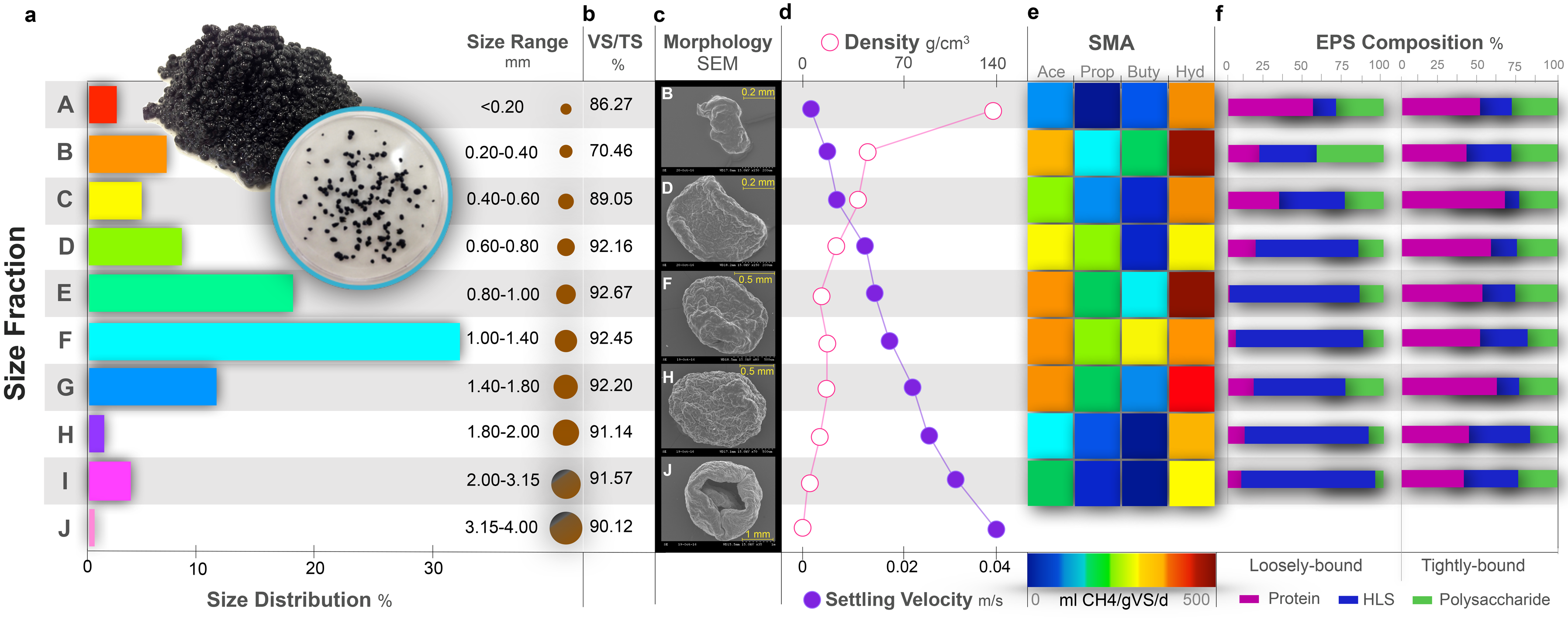
Physico-chemical and physiological data from granule size fractions, A – J. **(a)** Bar plot indicating the size ranges of respective fractions, along with relative volumetric contributions to the sludge; **(b)** VS proportions of TS; **(c)** typical scanning electron microscopy (SEM) micrographs of selected granules (from fractions B, D, F, H and J); **(d)** scatter plot illustrating density, and settling velocity, of granules (*n*=10) from each size fraction; **(e)** heat map depicting specific methanogenic activity (SMA) of sludge samples (*n*=3) from each size fraction (except fraction J) against acetate (Ace), propionate (Prop), butyrate (Buty) and H_2_/CO_2_ (Hyd); and **(f)** stacked bar charts showing relative concentrations of proteins, humic-like substances (HLS) and polysaccharides components in loosely-bound and tightly-bound-EPS extracted from each size fraction (except fraction J).

#### Granule ultrastructure

A clear gradient in aggregate ultrastructure was apparent across the size fractions (Fig S1). The smallest biofilms (Fractions A – C) presented as ‘flakes’; medium-sized granules (Fractions D – F) appeared better defined but were ‘flat’ (i.e. not spherical); larger granules (Fractions G – I) were distinctly spherical and more ‘granular’; and in the largest granules (Fraction J), large cracks and void spaces were apparent, and the granules appeared to have broken apart, losing structural integrity (Fig. 1).

#### Density and settleability

The density and settling velocity of granules were broadly linear across the size fractions (Fig. 1), but were inversely related: smaller granules (Fraction A) had high densities (104.6 g/cm^3^) but low settling velocities (average, 0.001 m/s), whilst large granules (Fraction J) were less dense (6.82 g/cm^3^) and more settleable (average, 0.041 m/s).

#### SMA across sizes

Granules from across all sizes were more methanogenically active against hydrogen than against the volatile fatty acid substrates tested (Fig. 1). Larger granules (Fractions H and I) were less active than smaller granules, regardless of substrate. Medium-sized granules (Fractions E – G) were generally the most active against all substrates.

#### EPS composition across granule sizes

There was no change in composition of tightly-bound (TB)-EPS across the size fractions - the proportion of each of the three examined components (protein, 53.2% (σ^2^, 7.1); humic-like substances (HLS), 23.0% (σ^2^, 7.5); and polysaccharides, the remaining 23.7% (σ^2^, 2.8)) was relatively stable. However, a gradient was apparent across the size fractions in the proportions of loosely-bound (LB)-EPS components; for example, the proportions of proteins and polysaccharides were high, and HLS were low, in small granules, whilst the obverse was the case for large granules (Fig. 1).

#### Microbial composition and diversity across size fractions

Amplicon sequencing data analysis for the ten separate size fractions (*n* = 30 samples) resulted in, from each fraction: 2 927 OTUs from an average of 71 772 ± 18 691 initial paired-end reads, 71 345 ± 18 596 reads prior to quality trimming with Sickle, and 53 239 ± 14 453 paired-end reads capable of being overlapped using PandaSeq.

Alpha diversity analysis indicated a strong, linear diversity gradient across the size fractions with significantly (p = 0.00019) higher rarefied richness in the smaller granules (Fraction A) than in the larger granules (Fraction J), and with significant differences between nearly every combination of fractions. Similarly, for Shannon entropy, small granules housed much more diverse communities than larger granules (p = 0.00012 between Fractions A and J), and there were significant differences between nearly every possible combination of fractions. Beta diversity analysis revealed a highly significant (p = 0.001) differentiation pattern across Fractions A to J using the Bray-Curtis (p = 0.001), unweighted UniFrac (Fig. S2), and the weighted UniFrac distance metrics (Fig. 2; p = 0.001).

**Figure 2.**
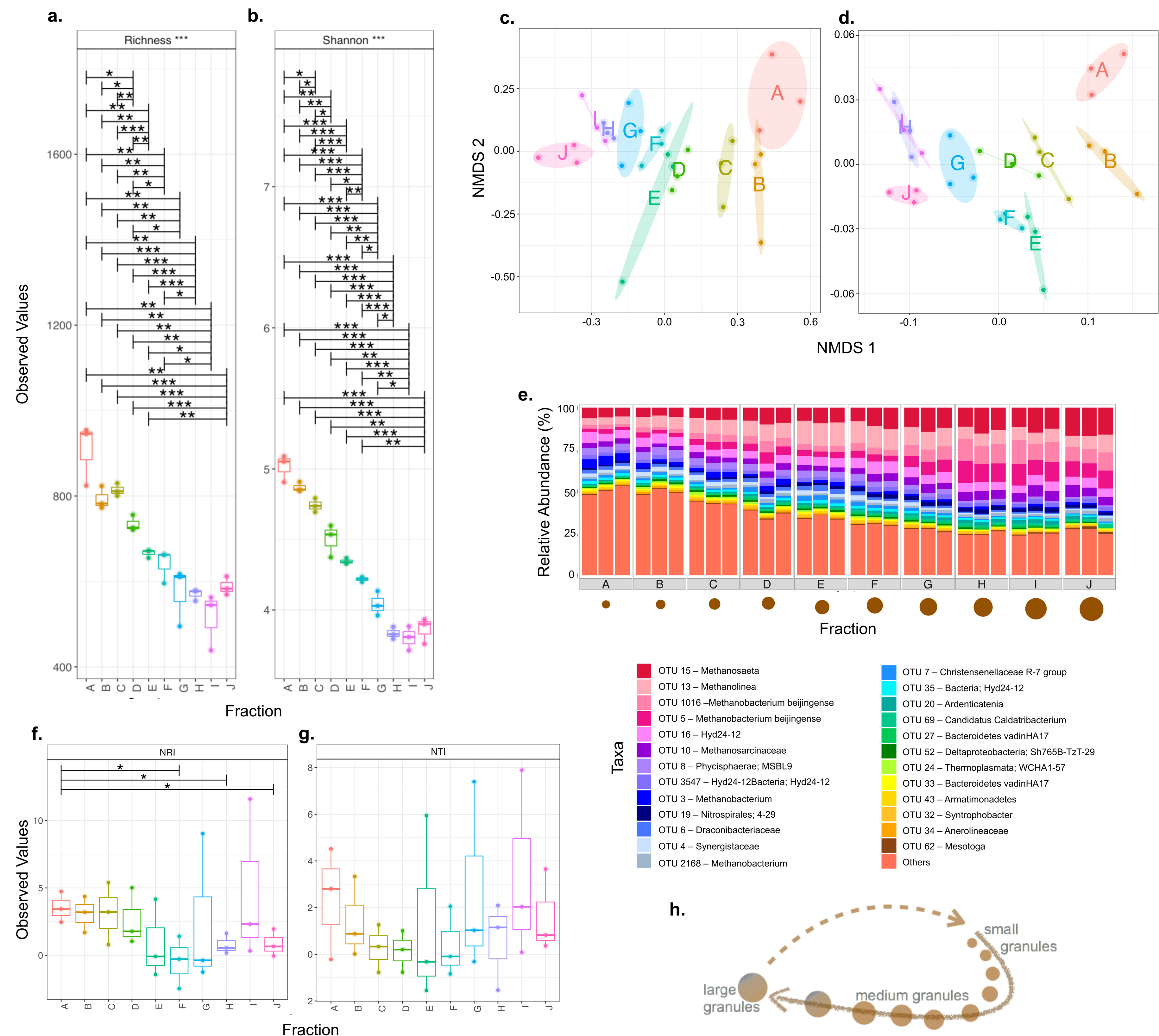
Microbial diversity, and community structure, in samples (*n*=3) from across each of the ten size fractions, A-J, according to variances in the 16S rRNA gene. *Alpha diversity*: box plot of the **(a)** rarefied species richness and **(b)** Shannon Entropy. *Beta diversity*: Non-Metric Multidimensional Scaling (NMDS) using **(c)** Bray-Curtis dissimilarity and **(d)** weighted UniFrac distances, where each point corresponds to the community structure of one sample, size fractions are indicated by colour, and the ellipses are drawn at a 95% CI; **(e)** community structure based on relative abundance of the top-25 most abundant OTUs from across each size fraction, where ‘others’ refers to all OTUs not included in the ‘top-25’; *Environmental Filtering:* **(f)** Net Relatedness Index (NRI) and **(g)** Nearest Taxa Index (NTI) calculated using the phylogenetic tree with presence/absence abundance; **(h)** depiction of the proposed granule growth trajectory, where the dashed line shows the proposed, closed life cycle. Lines for figures a, b, f & g connect two categories where the differences were significant (ANOVA) with * (P < 0.05), ** (P < 0.01), or *** (p < 0.001).

Archaea from the most abundant taxa (Fig. 2) were dominated by the two acetoclastic methanogens from the genus *Methanosaeta*; hydrogenotrophic methanogens from the genera *Methanolinea* and *Methanobacterium;* and the metabolically-diverse methanogens from the family *Methanosarcinaceae.* The 10 most abundant organisms included an interesting and diverse bacterial population: *Hyd24-12,* a candidate phylum in the Fibrobacteres-Chlorobi-Bacteroidetes superphylum; bacteria from the uncultured *Phycisphaerae* lineage; and the highly diverse *Nitrospirales* order. A gradual gradient in the relative abundance of the top-25 most abundant OTUs was observed across the size fractions (A to J), where the top-25 constituted 49.45% of the community in Fraction A but 78.48% of the community in Fraction J. Interestingly, the four most abundant OTUs, which were relatively more abundant with granule size, were all methanogenic archaea: *Methanosaeta* (5.71 ± 0.40% to 16.56 ± 0.42%), *Methanolinea* (4.30 ± 0.07% to 8.36 ± 2.01%), and two distinct classifications of *Methanobacterium beijingense* (2.31 ± 0.42% to 9.34 ± 1.28%; and 2.2 ± 0.42% to 9.41 ± 1.15%). The relative abundance of those methanogens, as a group, increased from 14.53% in Fraction A to 43.67% in Fraction J – nearly half of the entire microbial community.

NRI and NTI analyses provided U-shaped gradients across the size fractions (Fig 2). The smallest and largest granules clustered together phylogenetically (positive values) and were therefore more influenced by environmental factors. Meanwhile, the medium-sized granules (Fractions E – G) has slightly negative NRI values – tending towards phylogenetic dispersion.

#### Discriminant OTUs across sizes

Despite originating from the same environmental conditions, the overall community structure was observed to be significantly different between Fractions A – J and sPLS analysis identified 155 discriminant OTUs responsible for the observed changes (Fig. 3). Fourteen of the discriminant OTUs were methanogens, whilst all others were bacteria. There were three distinct groupings according to abundance, and across sizes: 41.3% of the discriminant OTUs were ‘upregulated’ in the smallest granules, 9% in the medium size fractions, and 49.7% in the largest size fractions.

**Figure 3.**
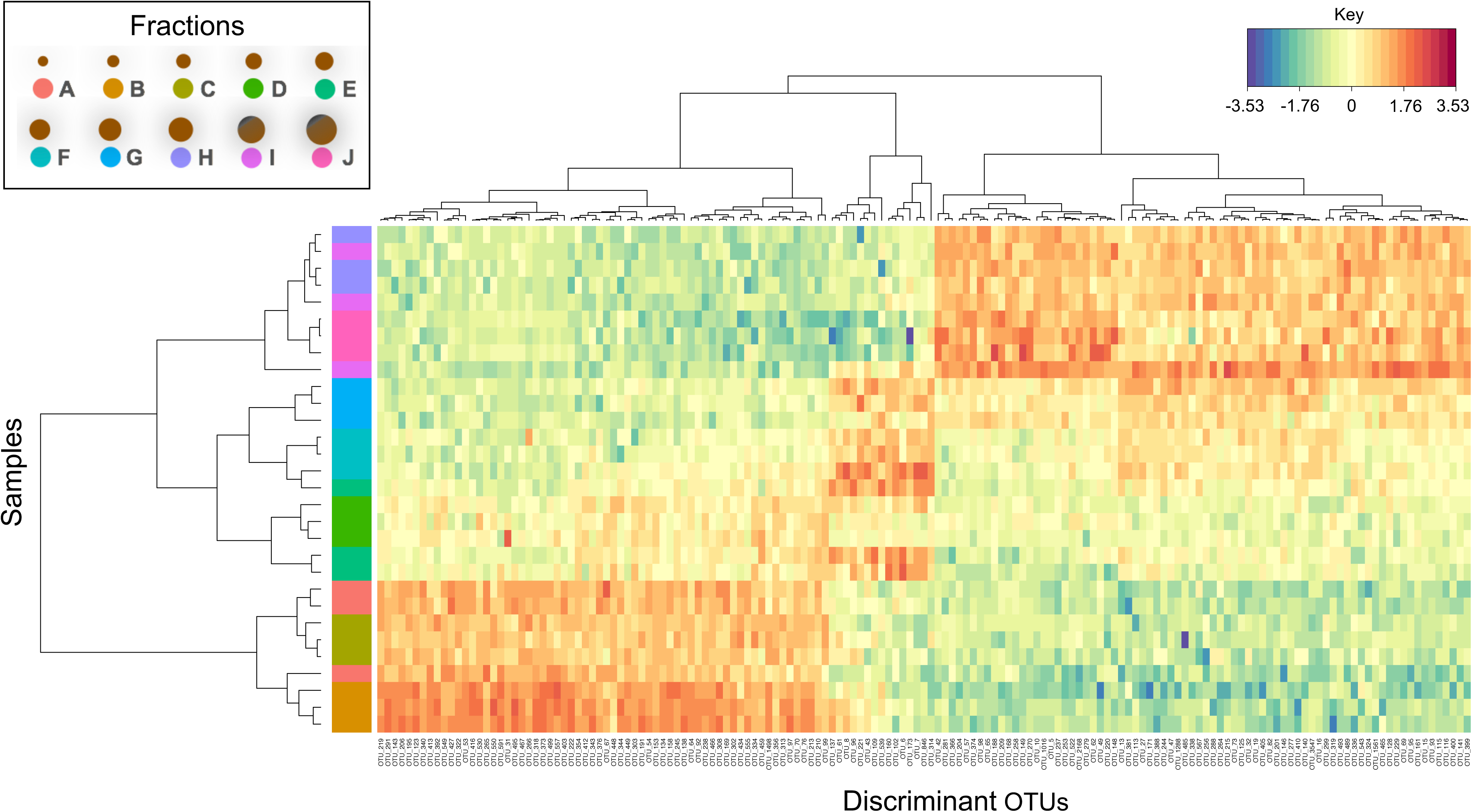
Heatmap of the discriminant OTUs across the ten size fractions (A – J; in triplicate, *n*=30) identified using sPLS-DA analysis with both rows and columns ordered using hierarchical (average linkage) clustering to identify blocks of OTUs of interest. Heatmap depicts TSS+CLR normalised abundances: high abundance (red) and low abundance (blue).

#### Correlations between physico-chemical differences and community structure

Strong positive and negative correlations were observed, based on granule size, between the 155 discriminant OTUs and the physico-chemical data (Fig. 4). The OTUs which were upregulated in the small granule fractions (A – C) showed strong positive correlations with density, and LB-EPS protein and polysaccharides. Discriminant OTUs in the medium-sized granules correlated significantly with SMA against acetate, propionate, butyrate and hydrogen – although, intriguingly, none of those OTUs were methanogenic archaea, indicating that various syntrophic and fatty-acid-oxidising bacteria may have been discriminant taxa in medium granules. There was a clear distinction in how OTUs upregulated in the medium-sized (Fractions D – G) and smaller granules correlated with meta-data. Correlations between upregulated OTUs in medium-sized granules and density, and settling velocity, were less significant than in small granules. The medium-sized granules appeared to represent the site of a transition zone, between small and larger granules, where the nature of such correlations shifted (from positive to negative, or *vice versa*). OTUs upregulated in the large granules (Fractions H – J) strongly positively correlated with settling velocity, and negatively correlated with SMA against hydrogen and butyrate.

**Figure 4.**
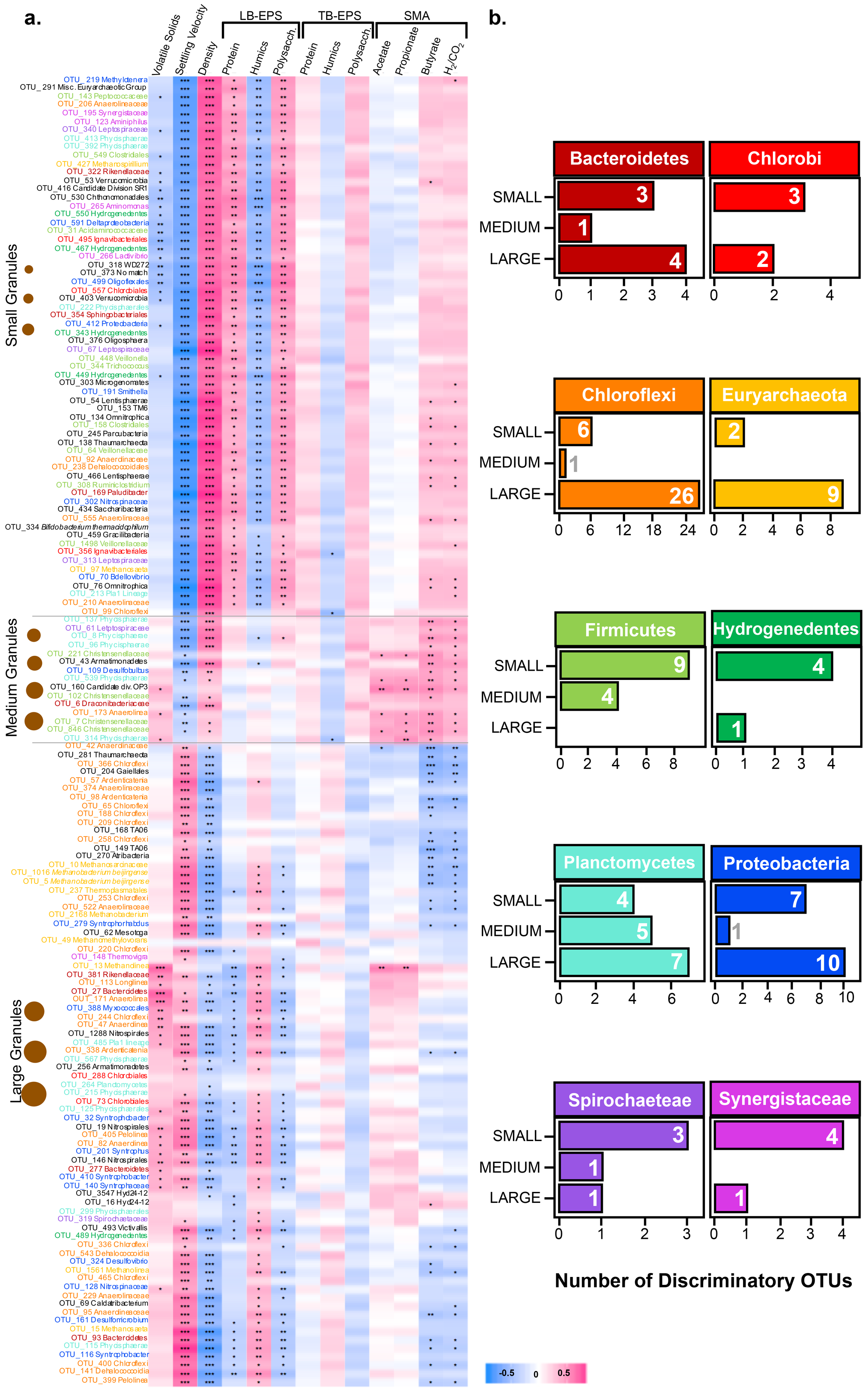
**(a)** Correlation plot depicting the 155 significantly discriminant OTUs coloured according to taxonomy (except where black), from sPLS analysis across the ten size fractions binned into three size groups: small (fractions A-C), medium (fractions D-G) and large (fractions H-J), and showing correlations with physico-chemical variables calculated using the Kendall rank correlation coefficient, where significant positive (pink) or negative (blue) correlations are marked with * (*Adj. P* < 0.05), ** (*Adj. P* < 0.01) or *** (*Adj. P* < 0.001); and **(b)** bar charts of the number of discriminant OTUs (x-axis) from major phyla that were found in small, medium, and large bins.

#### Euryarchaeota dominate

Although the community proportion of *Euryarcheota* increased with granule size, the entire group comprised only 69 OTUs in total, and was therefore not very diverse (Fig 5), which had the effect of reducing the total richness of the community in bigger granules (i.e. presumably, as granule size increases). This was tested by calculating the rarefied richness of the *Euryarcheota,* which was found to be fixed with granule size. In other words, although the rarefied richness fluctuated slightly, there was no trend toward reduced diversity within this phylum, which appeared to dominate as granules matured.

**Figure 5.**
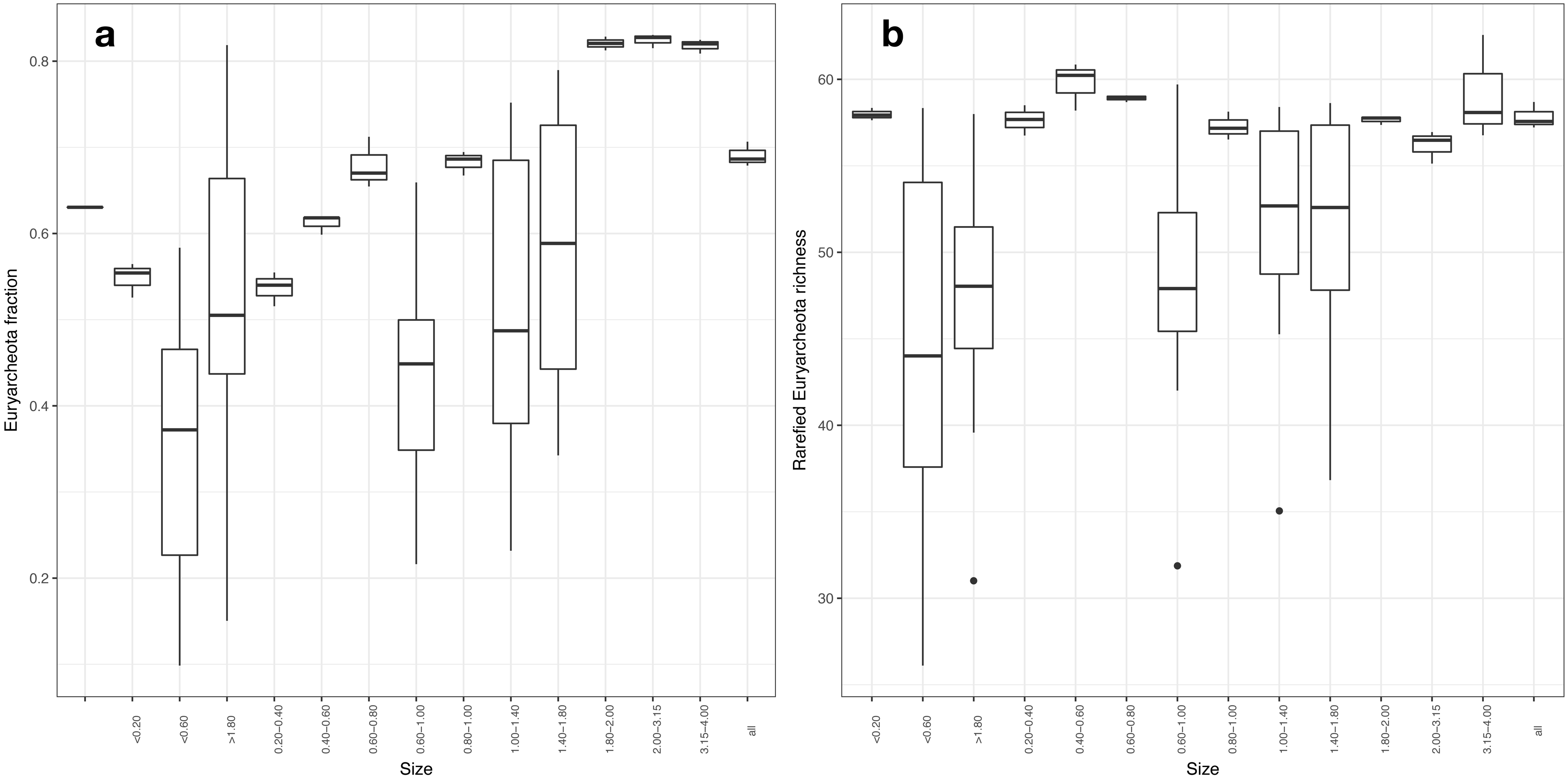
Plots showing that **(a)** as granules increase in size the fraction of *Euryarcheota* (which are 85% of methanogens) increases; and **(b)** rarefied *Euryarcheota* richness remains fixed with granule size.

### Discussion

One of the primary objectives of this study was to intensively characterise anaerobic granules from across a series of discrete sizes to identify drivers of community assembly during biofilm maturation. We hypothesised that distinct granule sizes correspond to stages in a biofilm life-cycle, in which small granules are ‘young’ and larger ones are ‘old’. Across each parameter explored, there were significant differences between granules of different sizes. As far we are aware, no such study has been reported before.

Volatile solids comprised a smaller proportion of the biofilms in fractions A – C. One of the key theories on granulation is the ‘*spaghetti theory*’, which proposes that during the initial stages of biofilm formation cells attach to inorganic nuclei [30,31], which may have made up an inorganic core comprising a larger proportion of the solids in the smaller granules. As further cells attach, and the biofilm grows, the organic fraction would become more important in larger granules, which is supported by the VS data presented.

A distinct gradient was observed in the ultra-structural features across the size fractions. Small granules were flaky and undefined, whilst the largest granules were spherical and – in some cases – beginning to break apart. Gradients in density and settleability profiles were also observed across size, whereby smaller granules were much more dense than larger granules, but had much lower settling velocities. Diaz *et al.* [18] also applied SEM using granules they had sliced in half revealing cross-sections, observing that largest granules had major cracks, and void spaces, that were less apparent in smaller granules. It is possible that such differences, or changes, in the structure of the granules also affects density: as granules become larger they acquire more cracks, channels and void spaces due to gas diffusion from the biofilm interior, which in turn renders the biofilm less tightly packed and – consequently – less dense. Furthermore, previous studies have described stratification of the sludge bed in anaerobic digesters using granules, where larger granules occupy the bottom and smaller ones the top of the bed [32–34] – this is interesting if granules in a bioreactor are to be considered as a meta-community or meta-organism. More specifically, however, avoiding biomass wash-out is a key consideration in applying anaerobic granules in bioreactors, and the findings lead us to conclude that settleability, rather than density, is the driving force for stratification.

There appears to have been a clear, linear gradient characterised by reducing diversity and converging community structure across the size fractions, from small to large, which was presumably across community assembly and biofilm maturation. This was somewhat counter to our initial assumption, which was that as the biofilm assembles and matures, diversity and rarefied richness would increase, especially as the biofilm simply contained significantly more cells. Both rarefied richness and diversity (measured by Shannon entropy) decreased significantly with granule size, due to the gradual dominance of a sub-group of the OTUs. Interestingly, the dominant, core group appeared to be comprised of four methanogenic archaea: *Methanosaeta, Methanolinea*, and two classifications of *Methanobacterium beijingense*, which are mainly hydrogenotrophic methanogens and may explain the high methanogenic activity measured against hydrogen.

*Methanosarcinaceae* were also present, which are able to metabolise a wide range of substrates including methylated amines, methanol, H_2_/CO_2_, acetate, dimethyl sulfide, methanethiol and sometimes carbon monoxide [35]. Abundant bacteria included the candidate phylum *Hyd24-12*, which are globally distributed but commonly found in anaerobic digesters where they are likely key fermenters, producing acetate and H_2_ from sugars [36], and supporting the metabolisms of the methanogens and *Nitrospirales*, which are the predominant nitrite-oxidisers playing critical roles in the biogeochemical cycling of nitrogen [37].

Sacchrolytic fermenters comprised a vast majority of the discriminant OTUs, with only a few notable exceptions. Within the subgroup of discriminant OTUs upregulated in the small granules, three were known, or likely, parasites: bacteria from the TM6 lineage; the *Parcubacteria* and a genus of Proteobacteria; and *Bdellovibrio* [38,39]. In the group of discriminants upregulated in the medium-sized granules, notable taxa included *Desulfobulbus.* Of the *Desulfobulbus* species to have thus far been isolated, all come from anaerobic environments with one particular species, the syntroph *Desulfobulbus propionicus*, capable of propionate oxidation with a methanogenic partner [40,41]. Other members of the genus are only known to be primary fermenters. The subgroup of discriminant OTUs upregulated in the large granules contained many of the top-25 most-abundant OTUs – constituting the emergence of a core microbiome. This group additionally included several syntrophic bacteria, such as *Syntrophorhabdus*, *Methanomethylovorans, Syntrophobacter* and *Desulfomicrobium* among others. Those syntrophs generally are sulfate-reducers found in habitats ranging from marine sediments to anaerobic digesters [42,43]. Finally, the largest granules contained the majority of the discriminant OTUs classified as *Euryarchaeota*. One OTU, in particular, was identified as *Methanolinea*, hydrogenotrophic methanogens in the top-25 most-abundant OTUs, and which were significantly positively correlated with SMAs against acetate and propionate.

Three alternative explanations may address the nature of the community assembly, and decreasing diversity, evident in the granules from across the size fractions studied. The first is based on the *neutral explanation* i.e. that communities are a balance between immigration and extinction [44]. In that case, reduced microbial immigration would result in reduced diversity. The second is based on the number of *functional groups* present in samples from across the size fractions studied. The relative abundance of distinct methanogenic archaea appeared to have increased with granule size, but since that functional group was not very diverse richness would be curtailed. The final hypothesis is based on a *competition effect* whereby better competitors would dominate functional groups and lead to reduced diversity in granules from across the sizes.

Our analysis determined *Euryarchaeota* as the increasingly-dominating group along the gradient of granule sizes, from small to large, but that diversity in the group was low. This indicates that the decreasing richness observed was a phenomenon associated with changing proportions of functional groups rather than with reduced diversity inside groups. The basic neutral model ignores any functional differences between organisms and treats the community purely as a balance between immigration and extinction. What is more likely is that there are functional niches whose abundance can change over time according to conditions. Within those groups, neutrality may, indeed, operate adding additional ecological complexity. However, and due to the complexity of the biofilm formation and maturation, the neutral model alone cannot simply explain assembly or diversity. Indeed, granules may become more strictly anaerobic with increasing size, creating ideal conditions for methanogenic populations to expand.

The observations from this study – on gradients in ultrastructure, activity, EPS composition and community structure – culminate in the proposal of a life-cycle model for anaerobic granular biofilms. This model (Fig. 2) proposes that granules begin as very small, compact and structurally-irregular, yet diverse, agglomerations of cells. Such granules are considered to be ‘young’. As the biofilm ages, it grows into a medium-sized, highly-active, structurally-stable entity with a less-diverse community structure, selecting rather for a community capable of efficient methane generation i.e. a consortium completing the anaerobic digestion process without significant accumulation of intermediate by-products. We consider those granules to be ‘ripe’. Further aging weakens the granule structure, and cracks and voids form. Activity decreases, likely due to structural inefficiencies in mass transport of substrates from one trophic group to another, but the diversity continues to converge primarily toward a methanogenic consortium. The granules may then be considered as ‘mature’. It is probable then, that the granules eventually break apart, but that the small fragments are still comprised of an active consortium and form the basis for new ‘young’ granules. The only observation that does not fully support this hypothesis is the set of observed gradients in rarefied richness and diversity – the linearity of the gradients does not necessarily indicate a circular trajectory – leading to the question, *what exactly is the fate of a large granule?* And, what is happening between, in this study, fractions J and A? In particular, the source of the additional richness in Fraction A is unclear, but it is likely that the surrounding medium (of wastewater, in the case of bioreactors) will provide the necessary additional diversity to cultivate new young granules.

Interestingly, medium-sized granules contributed a volumetric majority to the biomass used for this experiment. Those were also the most methanogenically active granules and appeared to have the most ‘stable’ ultrastructure. This indicates that the medium-sized granules may be the most stable; least open to immigration; and most important for methane production. Additionally, our NRI and NTI analyses demonstrated that the community structure of medium-sized granules was least influenced by environmental stresses. Equally, however, the smallest and largest fractions clustered together with NRI and NTI analysis, indicating that both were more vulnerable to change and environmental influence – and supporting somewhat the idea that both occupy pivotal points of change on a potential biofilm life ‘cycle’.

Moreover, the alpha diversity analysis showed that medium-sized granules were perhaps optimally diverse – containing a community rich in methane-producing archaea, but not over-dominated by them. It is possible then, that the digester system may self-regulate to select for medium-sized granules as a type of optimal growth phase in which critical trophic groups are maintained, suggesting the use of sophisticated ecological survival strategies. Furthermore, this may point to potential management strategies for digester operation where systems are managed to promote the emergence and existence of medium-sized granules.

## Conclusions

In summary, ecophysiolological and physico-chemical gradients were apparent in methanogenic granules across the highly-resolved set of size fractions investigated, indicating that aggregate size matters for both structure and function. It appeared that, as such biofilms developed, the microbial community significantly lost diversity. We conclude that this was associated with low-diversity, functional groups – in in this case the *Euryarchaeota* – becoming more dominant, due to a niche functional effect as developing granules became more anaerobic. The methanogens comprised the majority of a core microbiome across the life-stages of anaerobic granules. Medium-sized granules may be optimal in terms of structure and function, and granules may follow a biofilm life-cycle that self-selects for mostly medium-sized granules in a meta-organism that also includes smaller and larger aggregates. Indeed, the idea that operating a digester toward further selecting for medium-sized granules might result in optimally efficient conversions and bioenergy production. Finally, such granules provide ideal playgrounds to study community assembly, expansion, and succession of complex biofilm microbiomes, as well as very-high-throughput studies to investigate the response of replicated, whole microbial communities to environmental parameters and change.

## Author Contributions

ACT, SC, UZI and GC, designed the study. ACT performed all of the physico-chemical characterisation with assistance from CM and SM. IB and GG collaborated on EPS measurements and characterisation. ACT prepared the sequencing libraries. UZI wrote the scripts for data analysis, which was conducted by ACT. CQ contributed to application of ecological theory. Results were interpreted by ACT, IB, GG, UZI and GC. ACT drafted the paper and CQ, UZI and GC revised the document. UZI and GC are joint corresponding authors. All authors approve the paper and agree for accountability of the work therein.

## Acknowledgements

The authors thank NVP Energy for providing anaerobic sludge granules. SC was supported by the Engineering and Physical Sciences Research Council, UK (EP/J00538X/1). CM was supported by Erasmus and by the University of Turin and NUI Galway. CQ was funded by an MRC fellowship MR/M50161X/1 as part of the CLoud Infrastructure for Microbial Genomics (CLIMB) consortium MR/L015080/1. UZI was funded by NERC IRF NE/L011956/1. GC, SM and ACT were supported by a European Research Council Starting Grant (3C-BIOTECH 261330) and by a Science Foundation Ireland Career Development Award to GC. ACT was further supported by a Thomas Crawford Hayes bursary from NUI Galway, and to visit IB and GG by a Short-Term Scientific Mission grant through the EU COST Action 1302.

---

**Figure S1.**
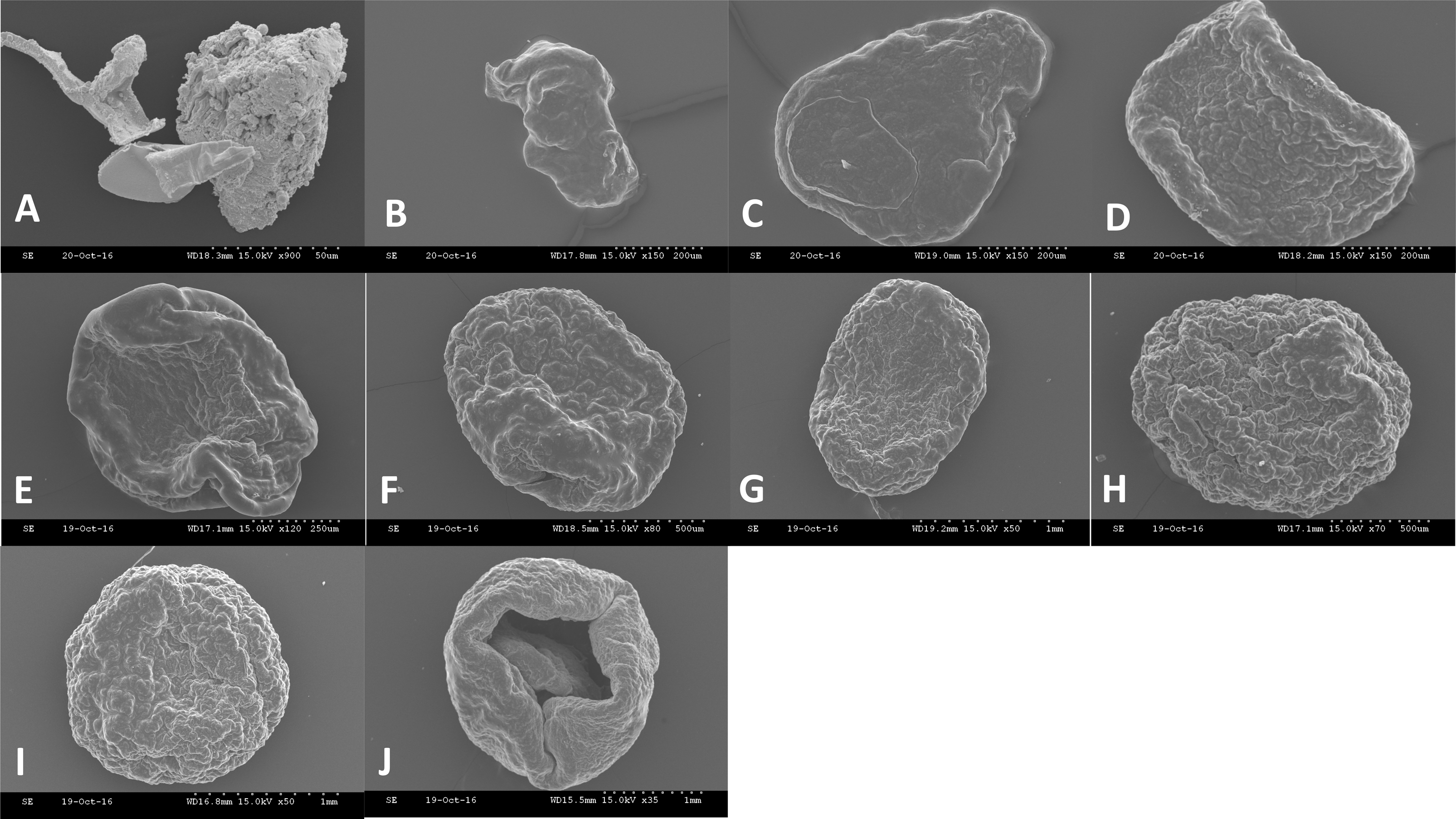
SEM micrographs of representative granules from size fractions A – J.

**Figure S2.**
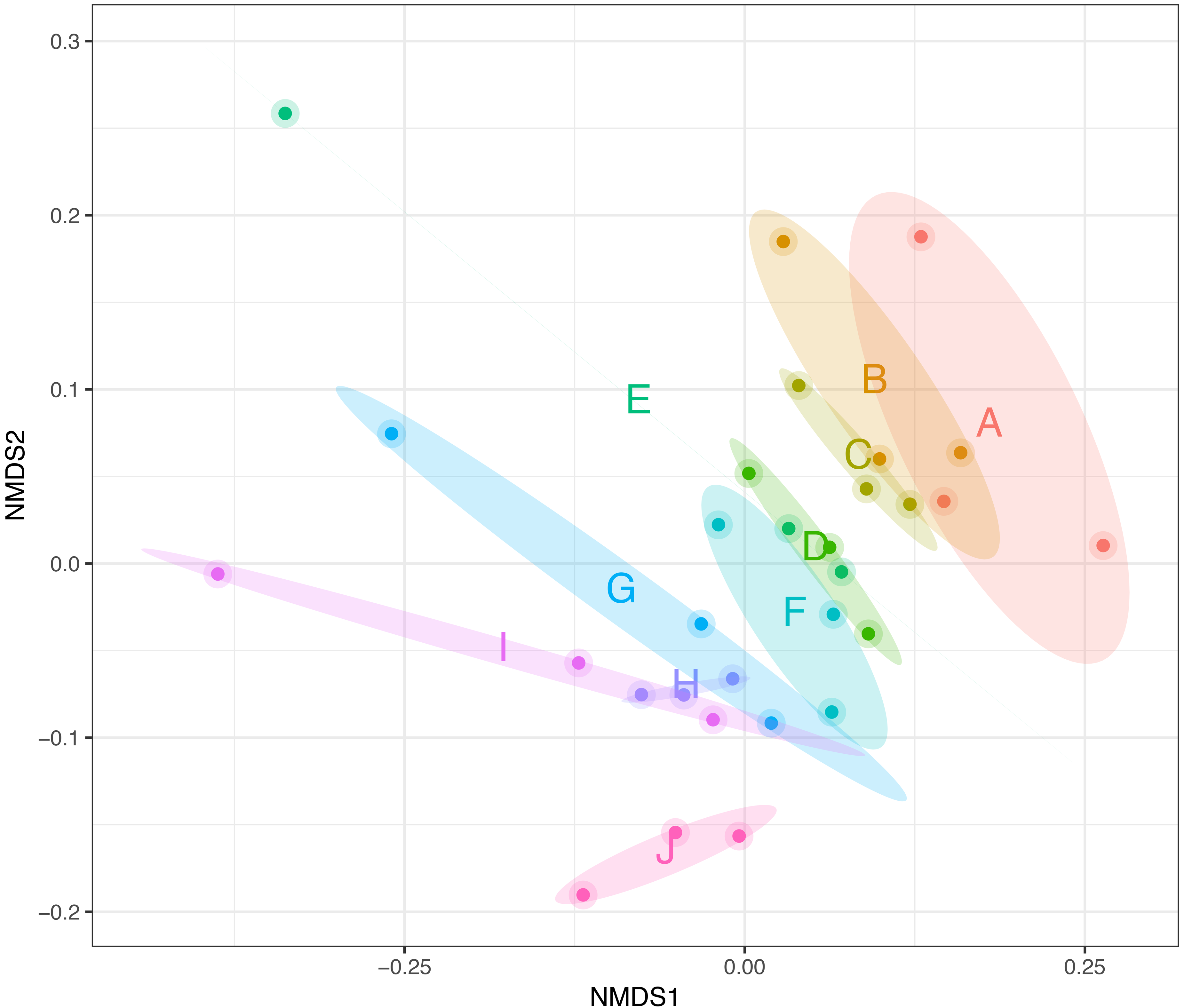
Non-Metric Multidimensional Scaling (NMDS) using UniFrac distances, where each point corresponds to the community structure of one sample, size fractions are indicated by colour, and the ellipses are drawn at a 95% CI.

## Supplementary Materials and Methods

### Bioinformatics

Sickle v1.200 (1) was used to trim and filter paired-end reads. This software used a sliding window technique and trimmed regions where the average base quality dropped below 20. Next, a length threshold of 10 bp was used to discard reads that fell below this length. BayesHammer (2) was applied from the Spades v2.5.0 assembler, which error-corrected the paired-end reads. Following this, pandaseq v(2.4) was used to assemble the forward and reverse reads into a single sequence spanning the entire V3-V4 region with a minimum overlap of 20 bp. This provided consensus sequences for each sample. Recent work (3,4) has shown that this pipeline significantly reduces substitution rates (which is the primary type of error encountered in datasets generated by the Illumina MiSeq platform).

Next, VSEARCH (v2.3.4) was used for OTU construction (the steps are documented at http://github.com/torognes/vsearch/wiki/VSEARCH-pipeline). First, all reads from each sample were pooled together while barcodes were added to keep track of from which sample the read originated. The reads were then de-replicated, sorted in order of decreasing abundance, and singletons were discarded. Next, the reads were clustered based on 97% similarity, followed by removing clusters with chimeric models built from more abundant reads (--uchime_denovo option in vsearch). To remove any chimeras that may have been missed, particularly in the case that they had parents that were absent from the reads or were present in very low abundance, a reference-based chimera filtering step (--uchime_ref option in vsearch) using a gold database (https://www.mothur.org/w/images/f/f1/Silva.gold.bacteria.zip) was applied. Finally, the OTU table was generated by matching the original barcoded reads against clean OTUs (a total of 2927 OTUs for n=30 samples) at 97% similarity (a proxy for species-level separation).

The assign_taxonomy.py script from the Qiime workflow (5) was used to taxonomically classify the representative OTUs against the SILVA SSU Ref NR database release v123 database. Phylogenetic distances between OTUs were resolved using kalign v2.0.4 (6) as a multisequence alignment (options –gpo 11 –gpe 0.85). Following this, FastTree v2.1.7 (7) generated the phylogenetic tree in NEWICK format and biome files for the OTUs were generated by combining the abundance table with taxonomy information using make_otu_table.py from the Qiime workflow.

### Statistical analyses

The vegan package (8) was used for alpha and beta diversity analyses. For alpha diversity measures we used: **(i)** *rarefied richness* – the estimated number of species/features in a rarefied sample (to minimum library size); **(ii)** *Shannon entropy* – a commonly used index to measure balance within a community; **(iii)** *Simpson index* – a measure of dominance that weighs towards the abundance of the most common OTUs and is less sensitive to rare(r) OTUs; **(iv)** *Pilou eveness*, which compares the actual diversity values to the maximum possible diversity value, and is constrained between 0 and 1.0, whereby lower values will indicate more variation in abundance between different OTUs in the community; and **(v)** *Fisher’s alpha* – a parametric index of diversity that assumes the abundance of OTUs following the log series distribution. Non-metric multidimensional scaling (NMDS) plots of OTUs using three different distance measures were made using Vegan’s metamds() function: **(1)** *Bray-Curtis*, which is a distance metric that considers only OTU abundance counts; **(2)** *Unweighted Unifrac*, which is a phylogenetic distance metric that calculates the distance between samples by taking the proportion of the sum of unshared branch lengths in the sum of all the branch lengths of the phylogenetic tree for the OTUs observed in two samples, and without taking into account their abundances; and **(3)** *Weighted Unifrac*, which is a phylogenetic distance metric combining phylogenetic distance with relative abundances. This places emphasis on dominant OTUs or taxa. Unifrac distances were calculated using the phyloseq package (9).

To understand multivariate homogeneity of group dispersions (variances) between multiple conditions, Vegan’s betadisper() function was used, in which the distances between objects and group centroids were handled by reducing the original distances (BrayCurtis, Unweighted Unifrac, or Weighted Unifrac) to principal coordinates and then performing ANOVA on them. Analysis of variance was performed using Vegan’s Adonis() against distance matrices (Bray-Curtis/Unweighted Unifrac/Weighted Unifrac). This function, refered to as PERMANOVA, fits linear models to distance matrices and used a permutation test with pseudo-F ratios.

Phylogenetic distances within each sample were further characterised by calculating the nearest taxa index (NTI) and net relatedness index (NRI). This analysis helped determine whether the community structure was stochastic (i.e. driven by competition among taxa) or deterministic (i.e. driven by strong environmental pressure). The NTI was calculated using mntd() and ses.mntd(), and the mean phylogenetic diversity (MPD) and NRI were calculated using mpd() and ses.mpd() functions from the picante package (10). NTI and NRI represent the negatives of the output from ses.mntd() and ses.mpd(), respectively. Additionally, they quantify the number of standard deviations that separate the observed values from the mean of the null distribution (999 randomisation using null.model-‘richness’ in the ses.mntd() and ses.mpd() functions and only considering taxa as either present or absent regardless of their relative abundance). The positive value of NTI indicates that species co-occur with more closely related species than expected by chance, with negative values suggesting otherwise. NTI measures tip-level divergences (putting more emphasis on terminal clades and is akin to “local” clustering) in phylogeny while NRI measures deeper divergences (akin to “global” clustering or “clumpedness”). For both NTI and NRI, values > +2 indicate strong environmental pressure, and values < −2 indicate strong competition among species as the driver of community structure. Based upon the recommendations given by Stegen *et al*. (11) we used only the top 1,000 most abundant OTUs for the calculations.

Sparse Projection to Latent Structure – Discriminant Analysis (sPLS-DA) was performed using the MixOmics package for R (12). This analysis constructed artificial latent components for predicted variables OTUs and response variables (categorical data matrix of different size fractions) by factoring these matrices into *scores* and *loading vectors* in a new space to achieve a maximum covariance between the scores of these two matrices. The loading vectors (with piece-wise coefficient for each OTU) were constructed so that the coefficients indicate the importance of each variable to define the component. Any non-zero coefficients for the loading vectors indicate genera that very significantly between the categories and are deemed discriminants. The OTU table was initially prefiltered by removing 1% of OTUs with low counts as based on author’s recommendation given at http://mixomics.org/mixmc/pre-processing/. After this step, we normalised the OTU table using Total Sum Scaling (TSS) on the OTUs followed by Centered Log Ratio (CLR), collectively referred to as TSS+CLR normalisation, and before applying the *splsda*() function. The *perf.plsda*() and *tune.splsda*() functions were initially used to predict the number of latent components (associated loading vectors) and the number of discriminants by initializing the *perf.plsda*() procedure with the total number components to be the number of size fractions considered in this study and then retaining the first two components as the classification error rates were minimum for these (<10%) using the centroid distance matrix in the procedure. The tune.splsda() function was then initialized with two components and using 5-fold cross-validation with 10 repeats (i.e. splitting the data into training and testing sets) identified the 155 discriminant OTUs amongst the size fraction. In both procedures we considered two metrics for classification error rates: the overall error rates and the balanced error rates (BER). Balanced error rates are measured between the predicted latent variables and the centroid of the class labels (size fractions considered in the study). BER accounts for differences in the number of samples between different size fractions. The OTUs that were deemed to be discriminants in the sPLS-DA analysis were selected and further analysed by determining correlations with external parameters (environmental meta data) using Kendall rank correlation with P-values adjusted for multiple comparisons using the Benjamini-Hochberg correction (13).

## References

1. Freeman C, Ostle N, Kang H. An enzymic &#39;latch&#39; on a global carbon store. Nature [Internet]. 2001 Jan 11;409:149. Available from: http://dx.doi.org/10.1038/35051650

2. Chapelle FH, O’Neill K, Bradley PM, Methé BA, Ciufo SA, Knobel LL, et al. A hydrogen-based subsurface microbial community dominated by methanogens. Nature [Internet]. 2002 Jan 17;415:312. Available from: http://dx.doi.org/10.1038/415312a

3. Martin W, Müller M. The hydrogen hypothesis for the first eukaryote. Nature [Internet]. 1998 Mar 5;392:37. Available from: http://dx.doi.org/10.1038/32096

4. Dolfing J, Larter SR, Head IM. Thermodynamic constraints on methanogenic crude oil biodegradation. Isme J [Internet]. 2007 Dec 13;2:442. Available from: http://dx.doi.org/10.1038/ismej.2007.111

5. Appels L, Lauwers J, Degrève J, Helsen L, Lievens B, Willems K, et al. Anaerobic digestion in global bio-energy production: Potential and research challenges. Renew Sustain Energy Rev [Internet]. 2011;15(9):4295–301. Available from: http://www.sciencedirect.com/science/article/pii/S1364032111003686

6. Seghezzo L, Zeeman G, van Lier JB, Hamelers HVM, Lettinga G. A review: The anaerobic treatment of sewage in UASB and EGSB reactors. Bioresour Technol [Internet]. 1998;65(3):175–90. Available from: http://www.sciencedirect.com/science/article/pii/S0960852498000467

7. van Lier JB, van der Zee FP, Frijters CTMJ, Ersahin ME. Celebrating 40 years anaerobic sludge bed reactors for industrial wastewater treatment. Rev Environ Sci Bio/Technology [Internet]. 2015;14(4):681–702. Available from: https://doi.org/10.1007/s11157-015-9375-5

8. Kato MT, Rebac S, Lettinga G. Anaerob c Treatment of of Low-Strength Brewery Wastewater in Expanded Granular Sludge Bed Reactor. Appl Biochem Biotechnol. 1998;76.

9. Lettinga G, van Velsen AFM, Hobma SW, de Zeeuw W, Klapwijk A. Use of the upflow sludge blanket (USB) reactor concept for biological wastewater treatment, especially for anaerobic treatment. Biotechnol Bioeng [Internet]. 1980 Sep 11;22(4):699–734. Available from: https://doi.org/10.1002/bit.260220402

10. Ir R, Lettinga G. Anaërobe zuivering van het afvalwater van de bietsuikerindustrie (2)*. 1976;0(2):1–6.

11. van Lier JB, Tilche A, Ahring BK, Macarie H, Moletta R, Dohanyos M, et al. New perspectives in anaerobic digestion. Water Sci Technol [Internet]. 2001 Jan 1;43(1):1–18. Available from: http://dx.doi.org/10.2166/wst.2001.0001

12. Hulshoff Pol LW, de Castro Lopes SI, Lettinga G, Lens PNL. Anaerobic sludge granulation. Water Res [Internet]. 2004;38(6):1376–89. Available from: http://www.sciencedirect.com/science/article/pii/S0043135403006705

13. Stoodley P, Sauer K, Davies DG, Costerton JW. Biofilms as Complex Differentiated Communities. Annu Rev Microbiol [Internet]. 2002 Oct 1;56(1):187–209. Available from: https://doi.org/10.1146/annurev.micro.56.012302.160705

14. Hall-Stoodley L, Costerton JW, Stoodley P. Bacterial biofilms: from the Natural environment to infectious diseases. Nat Rev Microbiol [Internet]. 2004 Feb 1;2:95. Available from: http://dx.doi.org/10.1038/nrmicro821

15. Datta MS, Sliwerska E, Gore J, Polz MF, Cordero OX. Microbial interactions lead to rapid micro-scale successions on model marine particles. Nat Commun [Internet]. 2016;7(May):1–7. Available from: http://dx.doi.org/10.1038/ncomms11965

16. Langenheder S, Székely AJ. Species sorting and neutral processes are both important during the initial assembly of bacterial communities. Isme J [Internet]. 2011 Jan 27;5:1086. Available from: http://dx.doi.org/10.1038/ismej.2010.207

17. Ahn Y. Physicochemical and microbial aspects of anaerobic granular biopellets. J Environ Sci Heal Part A [Internet]. 2000 Oct 1;35(9):1617–35. Available from: https://doi.org/10.1080/10934520009377059

18. Díaz EE, Stams AJM, Amils R, Sanz JL. Phenotypic properties and microbial diversity of methanogenic granules from a full-scale upflow anaerobic sludge bed reactor treating brewery wastewater. Appl Environ Microbiol. 2006;72(7):4942–9.

19. Trego AC, O’Sullivan S, Mills S, Porca E, Ijaz UZ, Collins G. Highly-replicated, whole-microbial communities in single anaerobic sludge granules respond reproducibly and distinctly to environmental cues. bioRxiv. 2018;

20. Rillig MC, Muller LAH, Lehmann A. Soil aggregates as massively concurrent evolutionary incubators. Isme J [Internet]. 2017 Apr 14;11:1943. Available from: http://dx.doi.org/10.1038/ismej.2017.56

21. APHA. Standard methods for the examination of water and wastewater. 21st ed. New York: American Public Health Association, Washington DC; 2005.

22. Frølund B, Palmgren R, Keiding K, Nielsen PH. Extraction of extracellular polymers from activated sludge using a cation exchange resin. Water Res. 1996;30(8):1749–58.

23. D’Abzac P, Bordas F, Van Hullebusch E, Lens PNL, Guibaud G. Extraction of extracellular polymeric substances (EPS) from anaerobic granular sludges: Comparison of chemical and physical extraction protocols. Appl Microbiol Biotechnol. 2010;85(5):1589–99.

24. Lowry, Oliver; Rosebrough, Nira; Farr, A. Lewis & Randall R. Protein Measurement With The Folin Phenol Reagent. J Biol Chem [Internet]. 1951;193(1):265–75. Available from: http://www.life.illinois.edu/biochem/355/articles/LowryJBC193_265.pdf

25. Frølund B, Griebe T, Nielsen PH. Enzymatic activity in the activated-sludge floc matrix. Appl Microbiol Biotechnol. 1995;43:755–61.

26. DuBois M, Gilles KA, Hamilton JK, Rebers PA, Smith F. Colorimetric Method for Determination of Sugars and Related Substances. Anal Chem [Internet]. 1956 Mar 1;28(3):350–6. Available from: https://doi.org/10.1021/ac60111a017

27. Colleran E, Concannon F, Golden T, Geoghegan F, Crumlish B, Killilea E, et al. Use of Methanogenic Activity Tests to Characterize Anaerobic Sludges, Screen for Anaerobic Biodegradability and Determine Toxicity Thresholds against Individual Anaerobic Trophic Groups and Species. Water Sci Technol [Internet]. 1992 Apr 1;25(7):31 LP–40. Available from: http://wst.iwaponline.com/content/25/7/31.abstract

28. Coates JD, Coughlan MF, Colleran E. Simple method for the measurement of the hydrogenotrophic methanogenic activity of anaerobic sludges. J Microbiol Methods [Internet]. 1996;26(3):237–46. Available from: http://www.sciencedirect.com/science/article/pii/0167701296009153

29. Griffiths RI, Whiteley AS, O’Donnell AG, Bailey MJ. Rapid Method for Coextraction of DNA and RNA from Natural Environments for Analysis of Ribosomal DNA- and rRNA-Based Microbial Community Composition. Appl Environ Microbiol [Internet]. 2000 Dec 5;66(12):5488–91. Available from: http://www.ncbi.nlm.nih.gov/pmc/articles/PMC92488/

30. Weignant W. The “spagetti theory” on anaerobic sludge formation, or the inevitability of granulation. In: Lettinga G, Sehnder A, Grotenhuis J, Hulshoff Pol L, editors. Granular anaerobic sludge: microbiology and technology. THe Netherlands: Pudoc. Wageningen; 1987. p. 146–52.

31. Jian C, Shi-yi L. Study on Mechanism of Anaerobic Sludge Granulation in UASB Reactors. Water Sci Technol [Internet]. 1993;28(7):171–8. Available from: http://wst.iwaponline.com/content/28/7/171

32. Yan Y-G, Tay J-H. Characterisation of the granulation process during UASB start-up. Water Res [Internet]. 1997;31(7):1573–80. Available from: http://www.sciencedirect.com/science/article/pii/S0043135496003545

33. Torkian A, Eqbali A, Hashemian SJ. The effect of organic loading rate on the performance of UASB reactor treating slaughterhouse effluent. Resour Conserv Recycl [Internet]. 2003;40(1):1–11. Available from: http://www.sciencedirect.com/science/article/pii/S0921344903000211

34. Connelly S, Shin SG, Dillon RJ, Ijaz UZ, Quince C, Sloan WT, et al. Bioreactor Scalability: Laboratory-Scale Bioreactor Design Influences Performance, Ecology, and Community Physiology in Expanded Granular Sludge Bed Bioreactors. Front Microbiol [Internet]. 2017;8:664. Available from: https://www.frontiersin.org/article/10.3389/fmicb.2017.00664

35. Oren A. The Family Methanosarcinaceae BT - The Prokaryotes: Other Major Lineages of Bacteria and The Archaea. In: Rosenberg E, DeLong EF, Lory S, Stackebrandt E, Thompson F, editors. Berlin, Heidelberg: Springer Berlin Heidelberg; 2014. p. 259–81. Available from: https://doi.org/10.1007/978-3-642-38954-2_408

36. Kirkegaard RH, Dueholm MS, McIlroy SJ, Nierychlo M, Karst SM, Albertsen M, et al. Genomic insights into members of the candidate phylum Hyd24-12 common in mesophilic anaerobic digesters. Isme J [Internet]. 2016 Apr 8;10:2352. Available from: http://dx.doi.org/10.1038/ismej.2016.43

37. Daims H. The Family Nitrospiraceae BT - The Prokaryotes: Other Major Lineages of Bacteria and The Archaea. In: Rosenberg E, DeLong EF, Lory S, Stackebrandt E, Thompson F, editors. Berlin, Heidelberg: Springer Berlin Heidelberg; 2014. p. 733–49. Available from: https://doi.org/10.1007/978-3-642-38954-2_126

38. Nelson W, Stegen J. The reduced genomes of Parcubacteria (OD1) contain signatures of a symbiotic lifestyle [Internet]. Vol. 6, Frontiers in Microbiology. 2015. p. 713. Available from: https://www.frontiersin.org/article/10.3389/fmicb.2015.00713

39. Yeoh YK, Sekiguchi Y, Parks DH, Hugenholtz P. Comparative Genomics of Candidate Phylum TM6 Suggests That Parasitism Is Widespread and Ancestral in This Lineage. Mol Biol Evol. 2016 Apr;33(4):915–27.

40. Pagani I, Lapidus A, Nolan M, Lucas S, Hammon N, Deshpande S, et al. Complete genome sequence of Desulfobulbus propionicus type strain (1pr3(T)). Stand Genomic Sci [Internet]. 2011 Mar 4;4(1):100–10. Available from: http://www.ncbi.nlm.nih.gov/pmc/articles/PMC3072085/

41. El Houari A, Ranchou-Peyruse M, Ranchou-Peyruse A, Dakdaki A, Guignard M, Idouhammou L, et al. Desulfobulbus oligotrophicus sp. nov., a sulfate-reducing and propionate-oxidizing bacterium isolated from a municipal anaerobic sewage sludge digester. Int J Syst Evol Microbiol [Internet]. 2017;67(2):275–81. Available from: http://ijs.microbiologyresearch.org/content/journal/ijsem/10.1099/ijsem.0.001615

42. Meyer B, Kuehl J V, Deutschbauer AM, Arkin AP, Stahl DA. Flexibility of Syntrophic Enzyme Systems in Desulfovibrio Species Ensures Their Adaptation Capability to Environmental Changes. J Bacteriol [Internet]. 2013 Nov 1;195(21):4900–14. Available from: http://jb.asm.org/content/195/21/4900.abstract

43. Copeland A, Spring S, Göker M, Schneider S, Lapidus A, Del Rio TG, et al. Complete genome sequence of Desulfomicrobium baculatum type strain (X(T)). Stand Genomic Sci [Internet]. 2009 Jul 20;1(1):29–37. Available from: http://www.ncbi.nlm.nih.gov/pmc/articles/PMC3035215/

44. Ofiţeru ID, Lunn M, Curtis TP, Wells GF, Criddle CS, Francis CA, et al. Combined niche and neutral effects in a microbial wastewater treatment community. Proc Natl Acad Sci [Internet]. 2010 Aug 31;107(35):15345 LP–15350. Available from: http://www.pnas.org/content/107/35/15345.abstract

## References

1. Joshi N, Fass J. Sickle: A sliding-window, adaptive, quality-based trimming tool for FastQ files (Version 1.33). 2011.

2. Nikolenko SI, Korobeynikov AI, Alekseyev MA. BayesHammer: Bayesian clustering for error correction in single-cell sequencing. BMC Genomics [Internet]. 2013 Jan;14(1):S7. Available from: https://doi.org/10.1186/1471-2164-14-S1-S7

3. Schirmer M, Ijaz UZ, D’Amore R, Hall N, Sloan WT, Quince C. Insight into biases and sequencing errors for amplicon sequencing with the Illumina MiSeq platform. Nucleic Acids Res. 2015;43(6).

4. D’Amore R, Ijaz UZ, Schirmer M, Kenny JG, Gregory R, Darby AC, et al. A comprehensive benchmarking study of protocols and sequencing platforms for 16S rRNA community profiling. BMC Genomics [Internet]. 2016 Jan;17(1):55. Available from: https://doi.org/10.1186/s12864-015-2194-9

5. Caporaso JG, Kuczynski J, Stombaugh J, Bittinger K, Bushman FD, Costello EK, et al. QIIME allows analysis of high-throughput community sequencing data. Nat Methods [Internet]. 2010 Apr 11;7:335. Available from: http://dx.doi.org/10.1038/nmeth.f.303

6. Lassmann T, Sonnhammer ELL. Kalign -- an accurate and fast multiple sequence alignment algorithm. BMC Bioinformatics [Internet]. 2005 Dec;6(1):298. Available from: https://doi.org/10.1186/1471-2105-6-298

7. Price MN, Dehal PS, Arkin AP. FastTree 2 – Approximately Maximum-Likelihood Trees for Large Alignments. PLoS One [Internet]. 2010 Mar 10;5(3):e9490. Available from: https://doi.org/10.1371/journal.pone.0009490

8. Oksanen J, Blanchet F, Kindt R, Legendre P, Minchin PR, O’hara R, et al. Vegan: community ecology package. R Package version 2.2–1. 2015.

9. McMurdie PJ, Holmes S. phyloseq: An R Package for Reproducible Interactive Analysis and Graphics of Microbiome Census Data. PLoS One [Internet]. 2013 Apr 22;8(4):e61217. Available from: https://doi.org/10.1371/journal.pone.0061217

10. Kembel SW, Cowan PD, Helmus MR, Cornwell WK, Morlon H, Ackerly DD, et al. Picante: R tools for integrating phylogenies and ecology. Bioinformatics [Internet]. 2010 Jun 1;26(11):1463–4. Available from: http://dx.doi.org/10.1093/bioinformatics/btq166

11. Stegen JC, Lin X, Konopka AE, Fredrickson JK. Stochastic and deterministic assembly processes in subsurface microbial communities. Isme J [Internet]. 2012 Mar 29;6:1653. Available from: http://dx.doi.org/10.1038/ismej.2012.22

12. Rohart F, Gautier B, Singh A, Lê Cao K-A. mixOmics: An R package for ‘omics feature selection and multiple data integration. PLOS Comput Biol [Internet]. 2017 Nov 3;13(11):e1005752. Available from: https://doi.org/10.1371/journal.pcbi.1005752

13. Benjamini Y, Hochberg Y. Controlling the False Discovery Rate: A Practical and Powerful Approach to Multiple Testing. J R Stat Soc Ser B [Internet]. 1995;57(1):289–300. Available from: http://www.jstor.org/stable/2346101

